# Stiffness Drives Endothelial Senescence and Inflammation in Aging and Doxorubicin-Induced Vascular Dysfunction

**DOI:** 10.64898/2026.01.30.702284

**Authors:** Camilla Romagnoli, Penelope M. Tsimbouri, Giuseppe Ciccone, Mariana Azevedo Gonzalez Oliva, Timo Zadegh Nazari Shafti, Massimo Vassalli, Delphine Gourdon

**Affiliations:** Centre for the Cellular Microenvironment (CeMi), University of Glasgow, Glasgow, UK; Institute for Bioengineering of Catalonia (IBEC), Barcelona, Spain; Baylor College of Medicine, Houston, TX, USA

**Keywords:** senescence, endothelium, stiffness, doxorubicin, aging

## Abstract

Age-related arterial stiffening affects over ∼ 60% of elderly individuals and is a major independent risk factor for cardiovascular mortality. Similarly, doxorubicin-induced cardiotoxicity affects up to ∼ 48% of cancer patients, limiting therapeutic options. Here, we demonstrate that vascular stiffness mechanically amplifies endothelial senescence phenotypes, identifying the extracellular matrix as a potential therapeutic target for both cardiovascular aging and chemotherapy-induced vascular dysfunction. Using polyacrylamide (PAAm) hydrogels to mimick soft (3.3 kPa) and physiological (30 kPa) arterial stiffness, and glass to reproduce pathological stiffening, we show that substrate rigidity enhances senescence markers including *β*-galactosidase activity, DNA damage, and inflammatory cytokine secretion in both therapy-induced and replicative senescence models. Critically, we identify a protective effect at physiological stiffness, where IL-6 and IL-8 secretion is minimized compared to both softer and stiffer conditions, suggesting an optimal mechanical therapeutic window. RNA sequencing reveals stiffness-dependent activation inflammatory pathways including chemotaxis and leukocyte migration. Our findings position vascular stiffening not just as a consequence but as driver of endothelial dysfunction, creating a positive feedback loop amenable to therapeutic intervention. These mechanobiological insights provide rationale for developing mechanical-based therapies in cardiovascular medicine and cardio-oncology, where targeting tissue mechanics alongside conventional approaches could improve clinical outcomes.

## Introduction

Over the past two centuries, average human life expectancy has more than doubled in the most advanced countries [1]. This demographic shift has brought age-related diseases such as cardiovascular disorders to the forefront, now representing the leading cause of death in older adults [2]. As the proportion of elderly individuals continues to rise, narrowing the gap between total and healthy lifespan has become an urgent biomedical challenge.

A key hallmark of aging is the accumulation of senescent cells (SCs) [3]. Senescence is a stable state of cell cycle arrest, first identified in fibroblasts by Hayflick and Moorehead [4]. It is typically triggered by persistent DNA damage, including telomere shortening, oxidative stress, oncogene activation, or genotoxic drugs, which activate the DNA damage response (DDR). DDR signaling converges on the tumour suppressor pathways p53/p21 and p16^INK4A^/pRB, enforcing irreversible proliferation arrest [5, 6].

Senescent cells display enlarged morphology, apoptosis resistance, cell cycle arrest, altered lysosomal activity, chromatin modifications, such as DNA foci, and they secrete a pro-inflammatory senescence-associated secretory phenotype (SASP) comprising cytokines, chemokines, and matrix metalloproteinases [7, 8]. Although several markers such as SA-*β*-Gal, *γ*-H2AX foci, p53 and SASP components are widely used, no single hallmark is exclusive to senescence, necessitating combined approaches for detection.

Endothelial senescence is observed in multiple organs, including in kidneys, brain and aorta, and endothelial cells (ECs) are among the earliest to undergo senescence in vivo [9, 10]. This process contributes to pathologies such as hypertension, diabetes, and dementia [11]. ECs are continuously exposed to biochemical stimuli (ROS, cytokines, cytotoxic drugs) and biomechanical forces (shear stress, stretch, and cell-cell contact), all of which can initiate DDR and trigger senescence [9]. Cancer therapies exacerbate this process: the anthracycline Doxorubicin (Doxo), although highly effective, is limited by cardiotoxicity affecting up to ∼ 48% of patients and manifesting as cardiomyopathy, heart failure, and vascular dysfunction [12]. It induces senescence in both cancer and non-cancer cells, and significant similarities have been observed between aged and Doxo-treated hearts [13, 14].

In parallel, aging is associated with changes in the extracellular matrix (ECM) that alter tissue mechanics and cellular behavior [15]. As vessels progressively stiffen with age, this creates an increasingly hostile mechanical environment that could accelerate endothelial dysfunction. While young healthy arteries maintain relatively low stiffness values (35–40 kPa [16, 17]), aging and disease can lead to substantial increases in vascular stiffness, approaching the GPa range [16]. This age-related vascular stiffening contributes to isolated systolic hypertension, left ventricular hypertrophy, and heart failure with preserved ejection fraction, conditions that collectively affect millions of patients worldwide and lack effective therapies [11].

While the biochemical mechanisms of vascular aging and Doxo toxicity are increasingly understood, the role of tissue mechanics in modulating these processes remains largely unexplored from a translational perspective. Cellular senescence is a key driver of both age-related vascular dysfunction and chemotherapy-induced cardiovascular toxicity [3, 5, 6], however, whether mechanical interventions could prevent or reverse senescence-associated vascular pathology remains unknown.

We hypothesized that substrate stiffness acts as a mechanical amplifier of endothelial senescence, potentially representing a target for cardiovascular therapeutics.

To test this, we developed an in vitro platform using polyacrylamide hydrogels spanning the physiological-to-pathological stiffness range. We selected a physiological stiffness of 30 kPa to represent healthy endothelium, a soft substrate of 3 kPa (one order of magnitude lower) to identify potential trends, and glass as a stiff substrate to mimic age-related endothelial stiffening. Given the wide range of reported endothelial stiffness values, we hypothesise that the selected conditions—3 kPa, 30 kPa, and glass—provide a comprehensive assessment of how stiffness influences senescence, and indeed, these choices align with previous research [18–20]. All the substrates are coated with type I bovine collagen, widely used for endothelial cell culture, in order to allow the correct adhesion of the cells to the substrates [21, 22].

These substrates were combined with two complementary senescence models: (1) Doxorubicin-induced senescence mimicking chemotherapy exposure, and (2) replicative senescence modeling natural aging. Our approach aimed to identify mechanically-regulated pathways amenable to therapeutic intervention in both cardiovascular aging and cardio-oncology.

## Results

### Doxorubicin treatment optimization

To establish a clinically relevant model of chemotherapy-induced endothelial dysfunction, we optimized Doxo treatment in HUVECs using a range of concentrations from 65 to 1000 nM [23–26]. A 24-hour treatment protocol was selected to mimic acute clinical exposure (Figure 1A).

**Figure 1.**
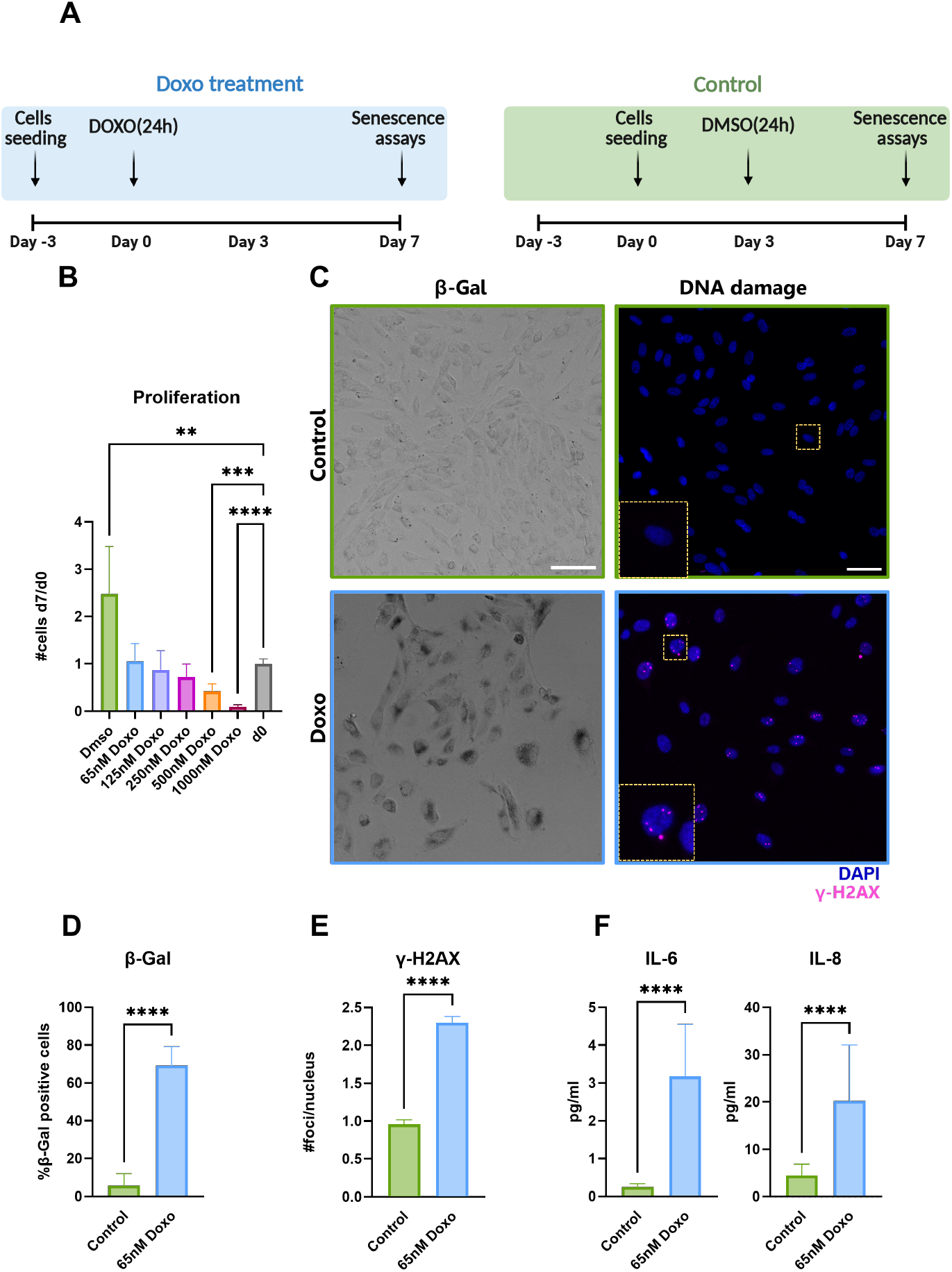
Doxorubicin dose optimization and senescence induction. (A) Schematic representation of the cell treatment protocol. For the Doxorubicin-treated (Doxo) samples (blue), cells were seeded at day -3, and Doxo was added at day 0 for 24h. For the control samples (green), cells were seeded at day 0, and treated with DMSO for 24h at day 3. For both conditions, the senescent assays were performed at day 7 after treatment. (B) Proliferation assay at day 7 following 24h Doxo treatment across doses from 65–1000 nM. Ratios represent cell number at day 7 relative to day 0 (mean *±* SD). (C) Left: representative bright-field images of the *β*-Galactosidase (*β*-Gal) staining at day 7, scale bar 100 *µ*m. Right: Representative immunofluorescence images: *γ*-H2AX (magenta), nuclei (blue); scale bar 20 *µ*m. (D) *β*-Gal positivity at day 7 (mean *±* SD). (E) DNA damage measured by *γ*-H2AX foci per nucleus (mean *±* SEM). (F) ELISA quantification of IL-6 and IL-8 secretion at day 7 (mean *±* SD). Statistical analysis: Mann-Whitney t-test; N = 2, n = 3. Significance: * ≤ 0.05, ** ≤ 0.01, *** ≤ 0.001, **** ≤ 0.0001.

Proliferation assays at day 7 identified three distinct response profiles: control cells showing normal growth (3-fold increase), high doses (500-1000 nM) causing cytotoxicity mimicking acute cardiotoxicity, and therapeutic doses (65-250 nM) inducing growth arrest without cell loss, modeling chronic vascular dysfunction (Figure 1B). The 65 nM concentration was selected as it induced robust senescence markers while maintaining cell viability (Figure S1), critical for studying long-term vascular complications.

The selected dose of Doxo (65 nM) induced classical senescence features: ∼ 70% *β*-galactosidase positivity (Figure 1D), 2.5-fold increase in DNA damage foci (Figure 1E), and elevated secretion of clinically relevant inflammatory mediators IL-6 (18-fold) and IL-8 (4.5-fold) (Figure 1F).

### Replicative senescence models natural vascular aging

To complement our therapy-induced model, we established replicative senescence through extended passaging, mimicking the natural aging process of the vasculature (Figure 2A). Late-passage cells (*>*P18) exhibited hallmarks consistent with aged human endothelium: proliferative arrest (Figure 2B), ∼ %50 *β*-galactosidase positivity (Figure 2D), doubled DNA damage (Figure 2E), and inflammatory cytokine secretion (Figure 2F).

**Figure 2.**
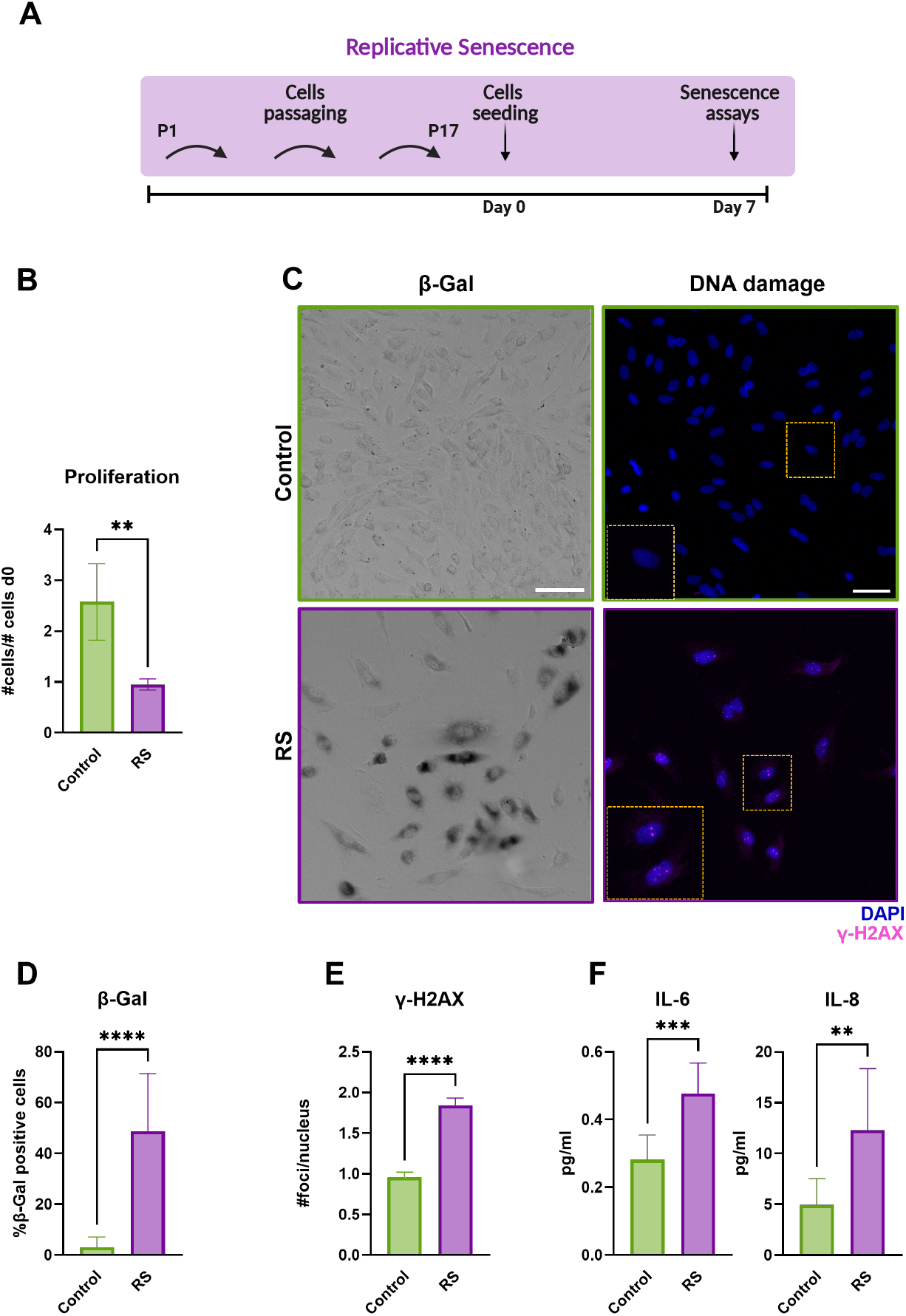
Replicative senescence induction. (A) Schematic representation of the protocol followed to obtain replicative senescence (RS). (B) Growth rate assessed as the ratio of cell number at day 7 to day 0 (mean *±* SD). (C) Left: representative bright-field images of the *β*-Galactosidase (*β*-Gal) staining at day 7, scale bar 100 *µ*m. Right: Representative immunofluorescence images: *γ*-H2AX (magenta), nuclei (blue); scale bar 20 *µ*m. (D) *β*-Gal positivity at day 7 (mean *±* SD). (E) DNA damage measured by *γ*-H2AX foci per nucleus (mean *±* SEM). (F) ELISA quantification of secreted IL-6 and IL-8 in RS cells versus controls. Statistical analysis: Mann–Whitney t-test. N=2–3, n=3. Significance: * ≤ 0.05, ** ≤ 0.01, *** ≤ 0.001, **** ≤ 0.0001.

### Stiffness amplifies the senescent phenotype

We tested the mechanical properties of the gels with nanoindentation. Performing force-displacement measure-ments, we measured the Young’s modulus (E) of the PAAm hydrogels and observed values of 3.3 *±* 0.3 kPa and 29.8 *±* 1.7 kPa, respectively for the soft and physiological substrates (Figure 3, A).

**Figure 3.**
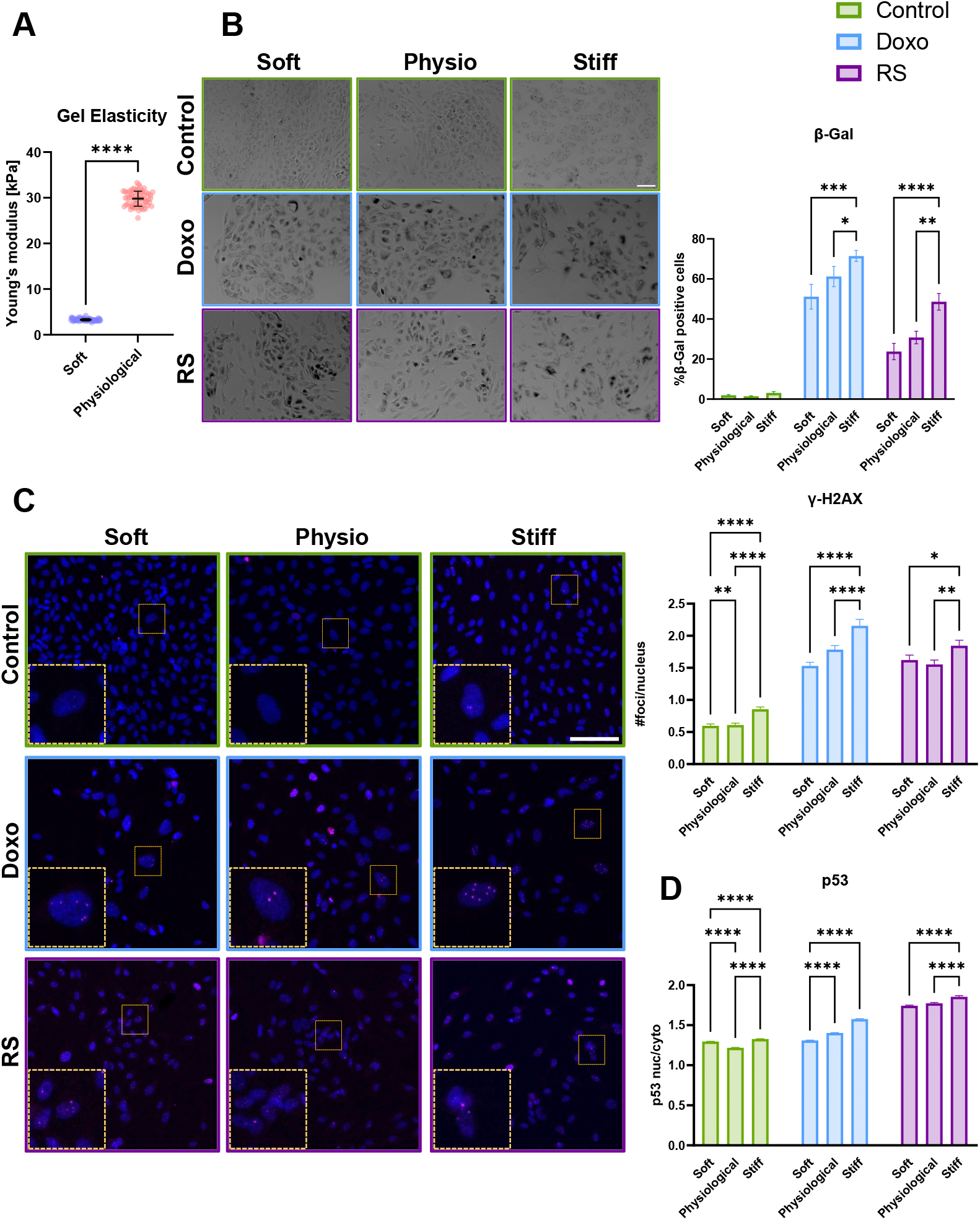
Impact of stiffness on the senescent phenotype. (A) Mechanical properties of PAAm hydrogels were measured by nanoindentation before collagen coating. Young’s modulus (mean *±* SD) was determined from force– displacement curves fitting via the Hertz model (see Results for values). Statistical analysis: Mann–Whitney test, number of gels tested N=2. (B) Percentage of *β*-Gal-positive cells relative to the total (mean *±* SD). Statistical analysis: two-way ANOVA with Tukey’s post hoc test; N=3, n=3 for control and Doxo; N=2, n=3 for RS. (C) DNA damage, assessed by *γ*-H2AX foci per nucleus (mean *±* SEM). Statistical analysis: Kruskal–Wallis test; N=3, n=3 for control and Doxo; N=2, n=3 for RS. (D) p53 nuclear localization, expressed as the nuclear-to-cytoplasmic mean intensity ratio (mean *±* SD). Statistical analysis: Kruskal–Wallis test; N=2, n=3. Significance: * ≤ 0.05, ** ≤ 0.01, *** ≤ 0.001, **** ≤ 0.0001.

To investigate the impact of substrate stiffness on the senescent phenotype, we analysed how key markers were influenced by different stiffness conditions. As previously reported, three substrates were included: soft (3 kPa), physiological (30 kPa), and stiff (glass, E ≈ GPa).

In order to assess the impact of substrates stiffness on the senescence phenotype, we quantified the number of *β*-Gal positively stained cells. Independently of substrate stiffness, the proportion of *β*-Gal-positive cells was higher in senescent populations compared to the control. We observed that cells cultured on glass displayed a higher number of *β*-Gal-positive cells compared to those on more compliant gels in both Doxo-induced and RS senescence models. However, substrate stiffness alone did not increase *β*-Gal staining in control cells (Figure 3, B).

We assessed DNA damage by quantifying *γ*-H2AX foci per nucleus. Regardless of substrate stiffness, both senescent populations tended to have an increased number of *γ*-H2AX foci per nucleus compared to the control, with RS cells showing higher foci counts than the Doxo-treated population (Figure 3, C). When analysing the response of the same cohort across different stiffnesses, for all the conditions, the number of foci per nucleus was significantly higher on glass compared to the softer substrates.

We examined p53 nuclear localization, a key marker of cell cycle arrest that was previously reported to increase in HUVECs after senescence induction [24, 27]. Across all conditions, RS cells displayed the highest nuclear p53 levels compared to control and Doxo-treated cells (Figure 3, D). When analysing substrate-dependent effects within each population, senescent populations exhibited a trend of increased p53 nuclear localization with increasing stiffness. For the control population, p53 nuclear localization showed a biphasic behavior with higher levels on the soft and stiff substrate compared to the physiological one.

### Stiffness amplifies the inflammatory phenotype

Cytokine secretion exhibited a stiffness-dependent, biphasic pattern across conditions. IL-6 and IL-8 levels were quantified in conditioned media collected 7 days after treatment or culture (Figure 4). For both cytokines, Doxo-treated cells secreted markedly higher levels than control and RS populations across all substrates. Importantly, for the Doxo-treated population, cytokines secretion was consistently the lowest on the physiological stiffness (30 kPa), whereas both soft and stiff substrates led to increased cytokine production. RS cells showed IL-6 and IL-8 levels comparable to controls across all stiffness conditions.

**Figure 4.**
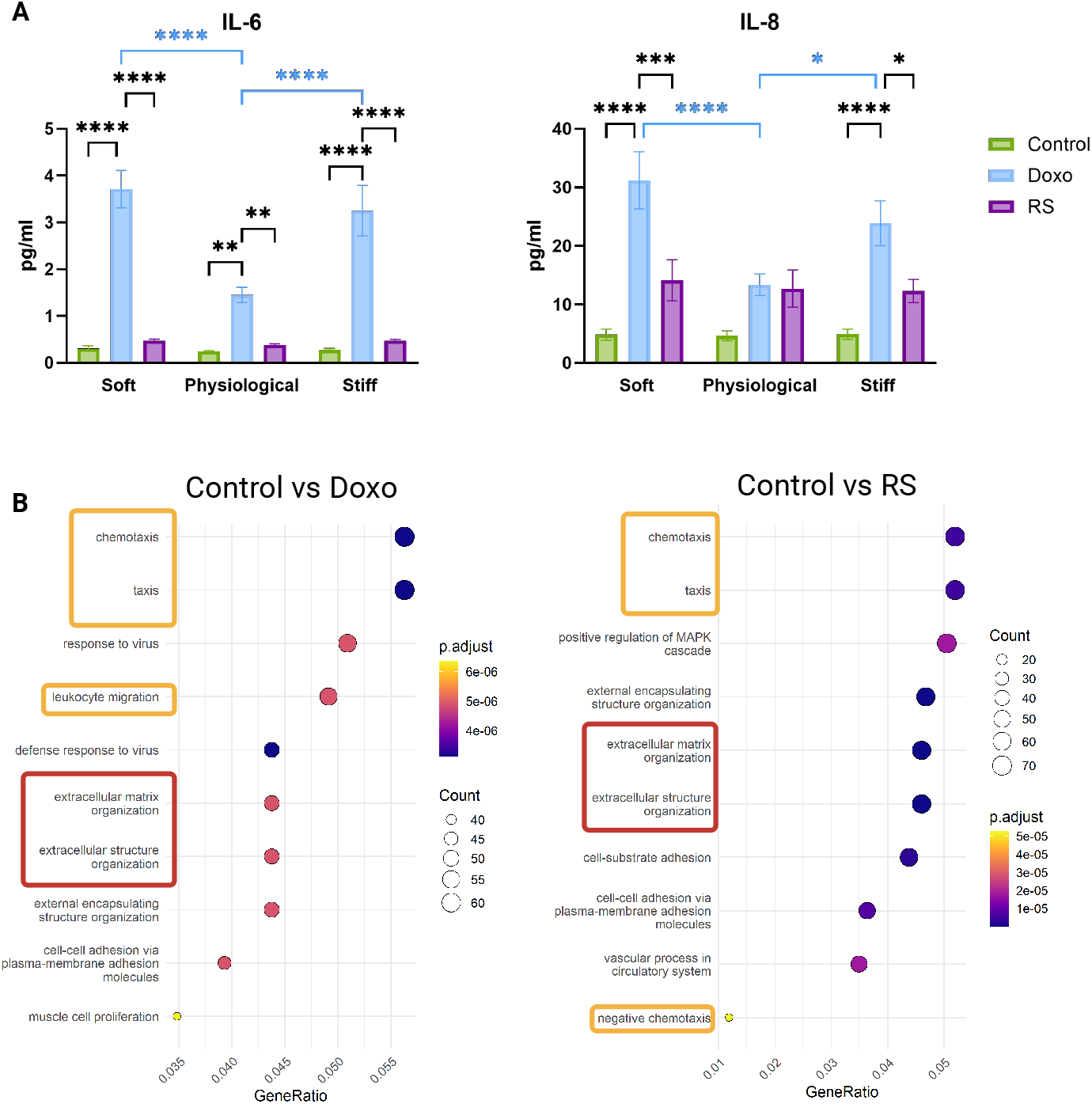
Effect of stiffness on the inflammatory phenotype. A) IL-6 and IL-8 secretion levels in control, Doxotreated, and RS populations after 7 days, quantified by ELISA and normalized to cell number (mean *±* SEM). Statistical analysis: two-way ANOVA with Tukey’s post hoc test (ELISA), N=3, n=3. Significance: * ≤ 0.05, ** ≤ 0.01, *** ≤ 0.001, **** ≤ 0.0001. B) Gene Ontology (GO) enrichment analysis displaying the top 10 Biological Process (BP) terms with highest statistical significance for Control vs Doxo and Control vs RS comparisons, respectively. For all GO term graphs, vertical axis shows GO terms and horizontal axis shows GeneRatio which corresponds to the genes of interest in the sample gene set divided over the total number of genes in the GO term. The size of each term dot represents the total count of genes identified in the sample gene set belonging to the term and the colour describes statistical significance according to the p-adjust legend. Recurring terms include chemotaxis, leukocyte migration, and extracellular matrix organization. Statistical analysis: DESeq2 (RNA-seq). N=3, n=3.

To evaluate whether these mechanical effects extended to the transcriptional level, we performed RNA sequencing on three independent pooled replicates per condition. GO enrichment analysis revealed recurring categories including chemotaxis and leukocyte migration, cell growth and division, and ECM organization (Figure S3). Comparisons performed on glass substrates highlighted enrichment of chemotaxis- and adhesion-related terms in both Doxo-treated and RS cells compared to controls. A stiffness-dependent pattern emerged in Doxo-treated samples: soft and stiff substrates were enriched for terms related to taxis and chemotaxis, whereas physiological stiffness was associated with categories linked to nuclear or cellular division (Figure S3). Differences in MAPK-related processes were also observed in the comparisons performed on glass.

Together, these data show that both cytokine secretion and transcriptional programs in senescent endothelial cells are strongly modulated by substrate stiffness, with distinct signatures between therapy-induced and replicative senescence.

### Substrate stiffness accentuates morphological and mechanical differences between senescent and non-senescent cells

To assess morphological changes, we quantified cell area after actin-phalloidin immunostaining. Both Doxo-treated and RS cells exhibited an enlarged area compared to controls across all stiffness conditions (Figure 5). On physiological and stiff substrates, RS cells tended to be larger than Doxo-treated cells.

**Figure 5.**
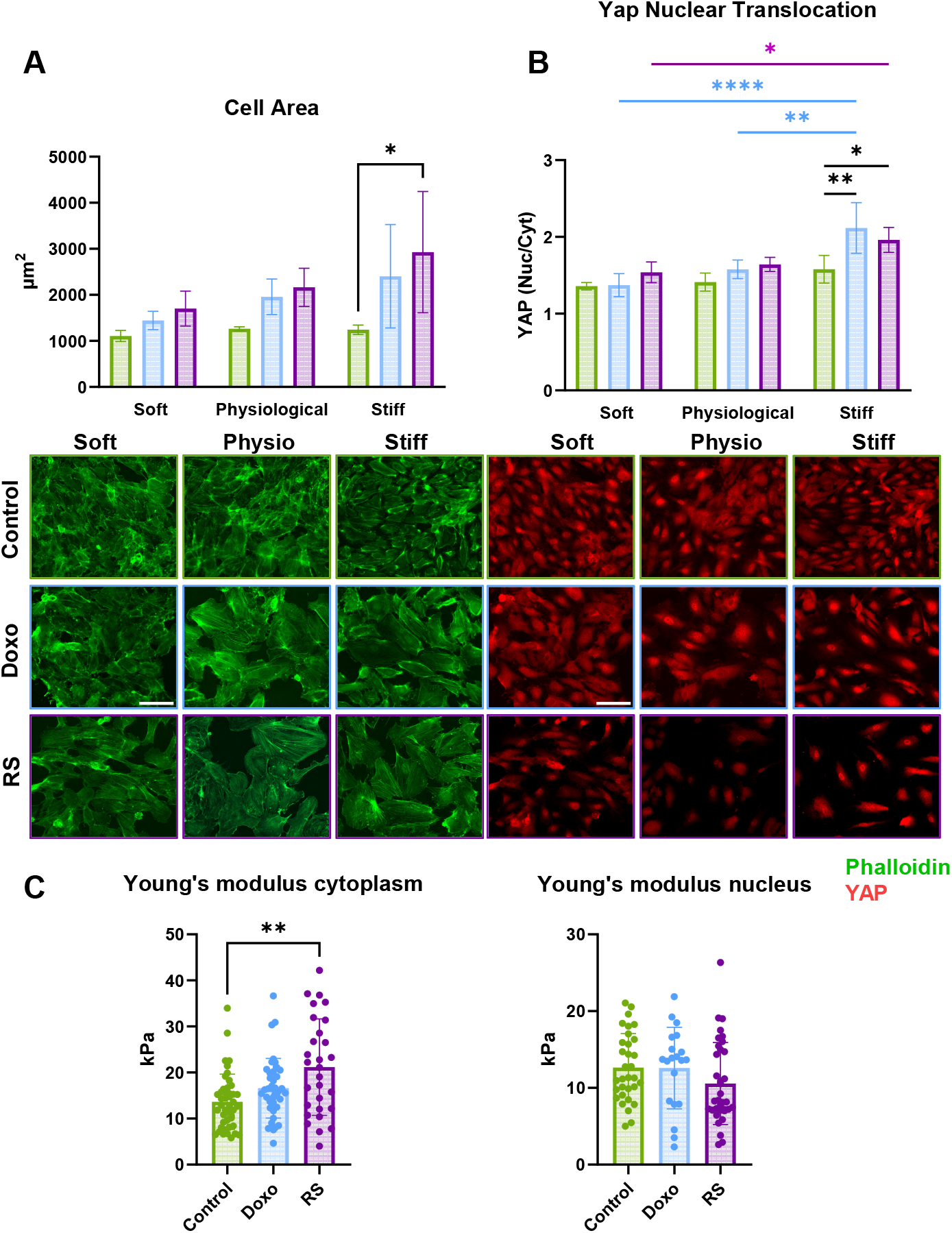
Substrate stiffness accentuates morphological and mechanical differences between senescent and non-senescent cells. (A) Cellular area quantified from actin-phalloidin immunostaining and image segmentation. Both Doxo-treated and RS cells exhibited a trend of increasing area compared to controls, with RS cells showing the largest values on stiffer substrates. Wide-field images acquired with a Zeiss Confocal 980 microscope (20× objective). Scale bar: 100 *µ*m. Mean *±* SD, *N* = 3, *n* = 3. Statistical analysis: two-way ANOVA with Tukey’s multiple comparisons test. (B) YAP nuclear translocation evaluated as nuclear-to-cytoplasmic intensity ratio. Images acquired with a Zeiss Confocal 980 microscope (20× objective). Scale bar: 100 *µ*m. Mean *±* SD, *N* = 3, *n* = 3. Statistical analysis: two-way ANOVA with Tukey’s multiple comparisons test. (C) AFM-based nanoindentation of cytoplasmic and nuclear regions. Mean *±* SD, *N* = 3. Statistical analysis: one-way ANOVA, Kruskal–Wallis test. Significance: * ≤ 0.05, ** ≤ 0.01, *** ≤ 0.001, **** ≤ 0.0001.

Since its discovery in 2011, the Yes-Associated Protein (YAP), a major component of the Hippo pathway, has become a widely recognised indicator of cellular perception and transmission of mechanical stimuli [28]. We evaluated YAP localization by measuring the nuclear-to-cytoplasmic intensity ratio. A stiffness-dependent trend in YAP nuclear translocation was observed across all conditions, and the values on glass resulted significantly higher than on the softer substrates for the senescent populations (Figure 5). Senescent cells displayed a trend of increasing nuclear YAP compared to controls on all substrates, with significant differences on glass.

Finally, we assessed intracellular mechanics using AFM-based nanoindentation. Cytoplasmic stiffness tended to higher values in senescent cells compared to controls, with RS cells showing the highest Young’s modulus values (Figure 5). Nuclear stiffness of senescent cells did not differ significantly compared to the control. These results indicate that substrate stiffness amplifies morphological, mechano-signaling, and mechanical distinctions between senescent and non-senescent cells.

## Discussion

### Substrate stiffness amplifies endothelial senescence

Our study reveals that substrate stiffness actively amplifies endothelial senescence, positioning vascular mechanics as both a driver of age-related dysfunction and a therapeutic target. The progressive increase in senescence markers from soft to stiff substrates, observed across *β*-gal activity, DNA damage, and p53 activation, demonstrates that mechanical forces universally modulate the senescent state.

The magnitude of mechanical amplification has direct clinical relevance. DNA damage foci nearly doubled on glass substrates even in control cells, suggesting that the increase in arterial stiffness observed during aging could significantly accelerate endothelial dysfunction [29, 30]. Moreover, pre-existing vascular stiffness, common in cancer patients with cardiovascular risk factors, could amplify chemotherapy-induced endothelial damage. Maintaining optimal vascular compliance may prevent senescence progression. Clinically, this suggests that interventions reducing arterial stiffness, such as angiotensin-converting enzymes (ACE) inhibitors, may indirectly attenuate senescence progression rather than merely treating symptoms.

### The homeostatic mechanical set point as a therapeutic target

Perhaps our most clinically relevant finding is the biphasic inflammatory response, with IL-6 and IL-8 secretion minimized at physiological stiffness ( ∼ 30 kPa) but elevated on both soft and stiff substrates. These results are consistent with previously reported data on smooth muscle cells [31], and we speculate that physiological stiffness represents an optimal mechanical environment that maintains endothelial homeostasis, whereas deviation in either direction triggers stress responses.

The mechanism likely involves optimal mechanotransduction at physiological stiffness. Our observation of intermediate YAP nuclear translocation at 30 kPa suggests balanced mechanosignaling, while deviations trigger stress responses. This is consistent with previous literature showing YAP/TAZ as mechano-rheostat [28]. The maintained YAP responsiveness in senescent endothelial cells contrasts with decreased YAP in aged stromal tissues [32], highlighting cell-type specificity. In our system, too little tension (soft substrates) may impair survival signaling through inadequate integrin clustering [33], while excessive tension (stiff substrates) triggers increasing engagement of the mechanotransductive molecular clutch and could potentially activate stress pathways, both converging on enhanced SASP.

This biphasic behavior for the Doxo-treated samples, is consistent with the GO results reporting terms related to chemokines and leukocyte migration on soft and stiff substrates and not on the physiological one.

From a therapeutic perspective, this biphasic response has several implications. First, it suggests that excessive softening of vessels could be detrimental and it identifies the existence of a homeostatic mechanical set point as a therapeutic target. Therapeutic interventions should aim to restore age-appropriate mechanics ranges rather than simply reduce stiffness. The dramatic reduction in inflammatory cytokines at optimal stiffness (*>* 50% decrease in doxorubicin-treated cells) suggests that mechanical normalization warrants investigation as a potential complement to anti-inflammatory approaches. Second, these results indicate that non-physiological stiffness, together with senescence induction, promotes leukocyte attraction, chemotaxis, and adhesion. This finding integrates previous observations showing that leukocyte migration is strongly influenced by substrate stiffness in bovine endothelial cells and that endothelial senescence enhances immune cell migration [34, 35]. Importantly, the combined effects of substrate stiffening, endothelial senescence, and leukocyte migration have been implicated in atherosclerotic plaque development [9]. Our data extend this framework by introducing chemotherapeutic-induced cardiotoxicity, underscoring the need to further investigate the cardiovascular effects of chemotherapeutic agents and positioning vascular stiffness as potential therapeutic target.

### Distinct mechanosensitive programs between Doxo-induced and replicative senescence

The differential responses between Doxo-induced and replicative senescence reveal distinct therapeutic opportunities. Doxorubicin-induced cells exhibited acute inflammatory signatures with IL-8 reaching 40 pg/ml on stiff substrates, nearly double that of replicative senescence. This distinction suggests different therapeutic strategies may be needed for age-related versus therapy-induced vascular dysfunction. These results mirror the rapid onset of clinical cardiotoxicity after Doxo treatment and suggests heightened mechanosensitivity following chemotherapy [24, 27]. For these patients, acute mechanical interventions during the peri-chemotherapy period could be particularly beneficial.

Conversely, replicative senescent cells showed stronger enrichment for ECM remodeling pathways, consistent with gradual matrix changes in vascular aging [36]. This identifies a different therapeutic strategy: long-term AGE breakers to prevent progressive stiffening. Notably, Doxycycline, which has MMP-inhibitory effects, reduces arterial stiffness in several experimental models, but its de-stiffening efficacy has not yet been demonstrated in human clinical trials [37].

### Breaking the pathological feedback loops

Our AFM measurements reporting increased cytoplasmic Young’s modulus (E), revealed that senescent cells themselves become stiffer, while RNA-seq showed upregulation of ECM remodeling genes. This could represent a pathological feedback loop: senescent cells stiffen and remodel their environment, which further amplifies senescence. This mechanism could explain the acceleration of vascular aging observed clinically, where pulse wave velocity increases exponentially rather than linearly with age [29]. Based on our findings and the existing literature, there are potential intervention points that warrant further investigation. For example, ACE inhibitors can reduce arterial stiffening, while ROCK inhibitors, like Netarsudil (approved in the United States for glaucoma and ocular hypertension) could reduce cellular contractility [38]. It is also widely known that exercise training reduces arterial stiffness by 15-20% [16]. Senescent cells can be targeted with senolytics, that could eliminate cells driving matrix remodeling [36], and finally YAP/TAZ inhibitors could prevent mechanical amplification [28, 39].

### Substrate stiffening as a hallmark of ageing

Our findings position substrate stiffening not merely as a consequence of aging, but as an active modifier of cellular ageing programs—qualifying it as a candidate hallmark of aging within the framework established by López-Otín and colleagues [3]. Hallmarks of ageing are defined as features that: (1) manifest during normal ageing, (2) worsen with accelerated ageing, and (3) when ameliorated, slow the aging process. Vascular stiffening satisfies the first two criteria unequivocally [29, 30], and emerging evidence suggests that interventions reducing arterial stiffness improve health-span [40]. Critically, our data demonstrate that stiffness acts as a gain control for senescence severity: the senescent triggers (doxorubicin or replicative exhaustion) produces more severe phenotypes on stiffer substrates. This amplification occurs through mechanotransduction pathways that are separable from the initiating insult, suggesting that targeting the mechanical microenvironment could ameliorate senescent cell burden independently of preventing senescence induction.

### Clinical biomarker development from mechanobiological insights

The stiffness-responsive SASP we identified offers potential novel biomarker strategies. The IL-6/IL-8 ratio differed dramatically between physiological and pathological stiffness, suggesting a mechanical inflammatory index combining cytokine levels with carotid-femoral PWV (cfPWV) could better predict cardiovascular risk than either alone. This is particularly relevant given that IL-6 and IL-8 are already validated cardiovascular biomarkers [41, 42]. Endothelial activation markers are under investigation (e.g., ANGPT2, soluble endoglin, endothelial microparticles), although these are not yet part of routine guidelines.

Our RNA-seq data revealed mechanically-regulated gene groups that could serve as circulating biomarkers. For instance, chemokine and leukocyte migration genes showed stiffness-dependent expression. Combining mechanical measurements such as cfPWV [43], with mechanically-responsive biomarkers could create powerful risk stratification tools.

### Conclusions: Toward mechanical medicine in vascular aging

This study establishes substrate stiffness as a critical amplifier of endothelial senescence. The discovery of a protective homeostatic mechanical set point, distinct mechanosensitive programs in different senescence types, and targetable feedback loops provides both mechanistic understanding and clinical opportunities. By demonstrating that vascular mechanics actively amplifies rather than passively accompanies cellular dysfunction, we provide rationale for incorporating mechanical diagnostics and therapeutics into cardiovascular medicine. As healthcare shifts toward precision approaches, our findings suggest that mechanical medicine, targeting the physical as well as biochemical disease environment, could significantly improve outcomes for patients affected by vascular aging and chemotherapy-induced cardiovascular toxicity [44].

### Limitations and future directions

Several limitations guide interpretation of our findings. Our 2D culture system lacks the 3D architecture and haemodynamic forces of blood vessels [45]. However, the consistency of mechanical effects across different senescence models suggests that our findings reflect fundamental mechanobiological principles. Future organ-on-chip studies incorporating flow and cyclic stretch would provide additional physiological relevance.

The polyacrylamide system provides purely elastic substrates, while it is increasingly recognised that vessels exhibit time-dependent responses, i.e., viscoelasticity, that impact tissue dynamics and functions. Recent work shows that substrate viscoelasticity affects cell fate [15], and future studies should examine how these time-dependent properties influence senescence. Additionally, while we identified correlations between mechanics and senescence, a deeper investigation must be conducted to decode the mechanism behind it, for this, perturbation experiments such as YAP knock down could help us confirm the key players mediating this mechanical amplification. Clinically, ACE therapies, that proved to have an impact on fibrosis and tissue remodelling, could be explored to observe their effect of senescence and inflammation.

## Methods

### Cell Culture

HUVECs (Caltag Medsystem) were cultured in endothelial cell growth medium (Promocell) supplemented with growth factors as recommended. Cells were incubated at 37 ^◦^C, 5% CO_2_, and passaged when confluent. After media removal, cells were washed twice in PBS and detached with trypsin for 3 min at 37 ^◦^C. Trypsin was neutralised with medium, cells were centrifuged for 5 min at 1300 rpm, and resuspended. Cell number was assessed with an automatic hemocytometer. Cells were seeded at 1:3 ratio for maintenance, and only passages *<*7 were used. Prior to seeding, flasks were coated with 1.5% bovine gelatin for 1 h at room temperature and rinsed with PBS.

### Doxorubicin-Induced Senescence

Doxorubicin (Abcam, ab120629) was dissolved in DMSO (25 mM), aliquoted, and stored at -80 ^◦^C. For working dilutions, a 100 *µ*M intermediate solution was stored at -20 ^◦^C for up to 2 weeks. Optimisation was performed with 65, 125, 250, 500, and 1000 nM; 500-1000 nM induced excessive cytotoxicity. Final experiments used 65-250 nM. DMSO controls were matched to the highest dose.

For senescence assays, cells were seeded at 2 × 10^4^ cells/cm^2^, treated for 24 h, rinsed twice with PBS, and maintained for 7 days with media changes every other day. A second medium exchange was performed 30-60 min post-treatment to remove residual drug. Control cells were seeded at 1 × 10^4^ cells/cm^2^ and exposed to vehicle for 24 h on day 3, ensuring equal culture duration. At day 7 post-treatment (or control seeding), senescence assays were performed (*β*-Gal, *γ*-H2AX, p53, cytokine ELISA). For mechanobiological studies, cells were seeded at 4 × 10^4^ cells/cm^2^, treated on day 4, and analysed 3 days later to maintain monolayer integrity.

### Proliferation Assays

Cells were seeded in 48-or 24-well plates (3 × 10^3^ or 10^4^ cells/well). NucBlue Live ReadyProbes (Thermo) were added 15 min prior to imaging (Zeiss Axio Observer, 5 × or EVOS M700, 10× ). Images were acquired on day 0, day 1, day 3, and day 7. For glass substrates, automated scanning covered ∼ 30% of each well. Nuclei were counted in Fiji using Otsu thresholding and the Analyze Particles function. Proliferation was expressed as nuclei count relative to *t*_0_.

### SA-*β*-Gal Assay

Cells were washed twice with PBS, partially fixed in 4% formaldehyde for 4 min, incubated with staining solution for 8 h at 37 ^◦^C, rinsed, and finally fixed for 10 min. Nuclei were counterstained with NucBlue. Imaging was performed with EVOS M700 at 20× . *β*-Gal positive cells were manually counted and normalised to DAPI-stained nuclei.

### ELISA

Conditioned media were collected on day 7, centrifuged (1000 g, 5 min), and stored at –80 ^◦^C. Media from 3 wells were pooled per condition. Human IL-6 and IL-8 DuoSet ELISA kits (R&D Systems) were used according to the manufacturer’s instructions. Optical density was read at 450 nm with 570 nm correction. Concentrations were interpolated from standard curves and normalised to cell number (nuclei count from DAPI-stained wells).

### Immunostaining

Cells were fixed with 4% paraformaldehyde (15 min), permeabilised with 0.1% Triton X-100 (15 min), and blocked with 1% BSA (1 h). Primary antibodies were incubated for 45 min at 37 ^◦^C, followed by secondary antibodies (45 min at 37 ^◦^C). Samples were mounted with DAPI antifade medium and stored at –20 ^◦^C. Reagents used for immunostaining: Formaldehyde (Fisher Scientific, 10630813, 4% in PBS); TritonX-100 (Sigma, X100, 0.1% in PBS); Bovine serum albumin (Sigma, A9418-50G, 1% in PBS); Vectashield Mounting media DAPI (Vector Laboratories, A300-081A, N/A); anti-*γ*-H2AX (Thermo Fisher, A300-081A, 1:800); anti-p21 (Proteintech, 67362-1-Ig, 1:100); anti-p53 (Proteintech, 60283-1-Ig, 1:100); anti-Yap (Santa Cruz, SC-101199, 1:200); anti-Tubulin (Proteintech, 66031-1-Ig, 1:100); anti-53BP1 (Novus Biotech, NB100-305, 1:2000); anti-*γ*-H2AX (Millipore, 05-636, 1:2000); Rabbit 647 Alexa (Life Technologies, A31573, 1:800); Mouse 488 Alexa (Life Technologies, A-11001, 1:800); Cy3 rabbit anti-mouse (Jackson Immuno, 315-165-300, 1:200); Alexa Fluor donkey 488 anti-rabbit (Thermo Fisher, A-21206, 1:200); 488 Phalloidin (Thermo Fisher, 6A12379, 1:250); CoraLite 594 Phalloidin (Proteintech, PF00003, 1:200); NucBlue Live Ready Probes (Thermo, R37605, 2 drops/mL).

### Image Analysis

Segmentation was performed with Cellpose [46], using the *cyto3* and *deblur cyto3* models. Channel configuration was adjusted per dataset (protein channel for cell body, DAPI for nuclei). Output masks were analysed in CellProfiler [47] to extract intensity, size, and shape features. Pipelines are available at GitHub repository of Camiroma .

### RNA Sequencing

RNA was extracted with RNeasy Micro Kit (Qiagen) and eluted in RNase-free water. DNase digestion was included. RNA quality and concentration were assessed by Nanodrop 2000. Libraries were prepared by Genewiz (Azenta) using the Illumina stranded mRNA protocol (polyA selection) and sequenced as 2 × 150 bp paired-end reads ( ∼ 20M/sample). Biological replicates were pooled to balance costs [48, 49]. We performed two main types of analysis: a graphic visualisation of the most represented genes and a Gene Ontology (GO) study of the most differentially expressed genes. Raw counts and TPM values were obtained. For the former, TPM *>*100 in at least one sample was used as threshold for visualization. This selection allowed us to visualise possible genes overrepresented in one sample and under-represented in other samples. For the Heatmap, we used the function *pheatmap* [50]. For the GO analysis, performed on raw counts, we used the DGElist function from the edgeR package [51] to create a digital gene expression, which allows for single replica statistical analysis. After normalising, we used the clusterProfiler package [52] on the org.Hs.eg.db database [53], and finally, we plotted the results using the ggplot2 package [54].

### Atomic Force Microscopy

Cells were seeded at 4 × 10^4^ cells/cm^2^ on collagen-coated glass-bottom dishes (100 *µ*g/mL). AFM (Nanosurf) was performed with PFQNM-LC-V2 probes (radius 70 nm, resonance frequency 80 Hz, *k* = 0.1 N/m). Force spectroscopy was performed in media, with indentation depth 2 *µ*m, speed 100 *µ*m/s, and 2000 data points. Young’s modulus was calculated using the Hertz model assuming *ν* = 0.5 [55].

### Polyacrylamide Hydrogels

Hydrogels were prepared as previously presented [56]. Coverslips were silanised with 3-(Acryloyloxy)propyltrimethoxysilane. Acrylamide/bisacrylamide mixtures (see Table 1) were polymerised between silanised coverslips and RainX-treated slides, then detached in milliQ water. Hydrogels were functionalised with sulfo-SANPAH (200 *µ*g/ml, UV 365 nm, 20 min) and coated with collagen type I (100 *µ*g/ml, 1 h). Mechanical properties were measured with a Chiaro Nanoindenter (Optics11) using a spherical probe (radius 27.5 *µ*m, *k* = 0.56 N/m). Young’s modulus was calculated from force–displacement curves using the Hertz model. Dynamic measurements (DMA) were performed with a 3 *µ*m probe at oscillation frequencies 1–10 Hz to determine storage (*E*^*′*^) and loss moduli (*E*^*′′*^).

**Table 1:** PAAm hydrogel recipes used in this study.

| Reagent ($\mu\text{L}$) | Soft | Physiological |
| --- | --- | --- |
| 40% Aam | 50 | 125 |
| 2% BisAam | 75 | 75 |
| mQ H<sub>2</sub>O | 336.7 | 261.7 |
| TEMED 1.5% | 33.3 | 33.3 |
| APS 10% | 5 | 5 |
| Tot Volume | 500 | 500 |

### Statistical Analysis

All analyses were performed in GraphPad Prism 10.4.1. Data are mean ± SD or mean ± SEM, as specified in each case. Normality was tested with D’Agostino-Pearson. Two-group comparisons: unpaired t-test (Welch’s correction if required) or Mann–Whitney test. Multiple-group comparisons: one-way/two-way ANOVA with Tukey or Kruskal–Wallis post hoc tests. *N* indicates experimental or biological replicates, whereas *n* indicate technical replicates. The formers refer to independent experiments performed on different days using separately prepared cell cultures, while technical replicates refer to repeated measurements within the same experiment, e.g., different coverslips. Significance: * ≤ 0.05, ** ≤ 0.01, *** ≤ 0.001, **** ≤ 0.0001.

## Acknowledgements

C.R. acknowledges funding from the Engineering and Physical Sciences Research Council (EPSRC) Scholarship under award EP/T517896/1 – 312561-05. D.G. acknowledges the Royal Society under the Wolfson award RSWF/FT/191020. We thank the members of the Centre for Cellular Microenvironment (CeMi), University of Glasgow, for their support. We thank the CRCHUM and members of the Rodier lab for welcoming us and sharing techniques and protocols. We thank Dr. Marco Cantini for feedback and suggestions.

## Notes

https://github.com/camiroma/thesis-cellprofiler-pipelines/tree/main

### Competing Interest Statement

The authors have declared no competing interest.

