## Supplementary for "Stiffness Drives Endothelial Senescence and Inflammation in Aging and Doxorubicin-Induced Vascular Dysfunction"

### Supplementary Results

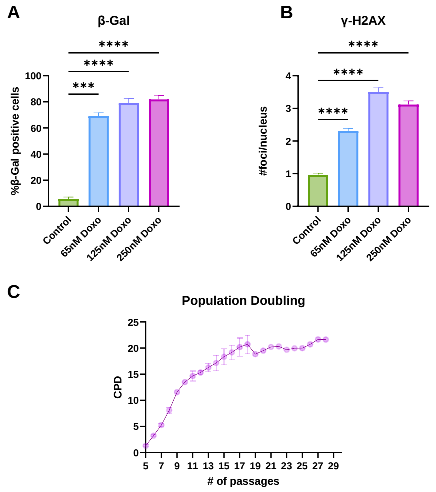


Figure S1: **Senescence induction** (A) $\beta$-galactosidase staining at day 7, expressed as % positive cells (mean ± SD). All selected doses (65, 125, 250 nM) showed 70% positivity. Statistical analysis: one-way ANOVA with Kruskal–Wallis post-test; N = 2, n = 3. (B) $\gamma$-H2AX foci per nucleus at day 7 (mean ± SEM), All selected doses (65, 125, 250 nM) showed presence of DNA damage. Statistical analysis: one-way ANOVA with Kruskal–Wallis post-test; N = 2, n = 3. (C) The Cumulative Population Doubling (CPD) level was calculated as $CPD = PDL_0 + \log(\frac{N_f}{N_i}) \times \log(2)^{-1}$. The graph reports the CPD as a function of time (days) of culture. Each point represents a passage, from P5 to passage P28 (mean ± SEM). N=1, n=1 or 3.

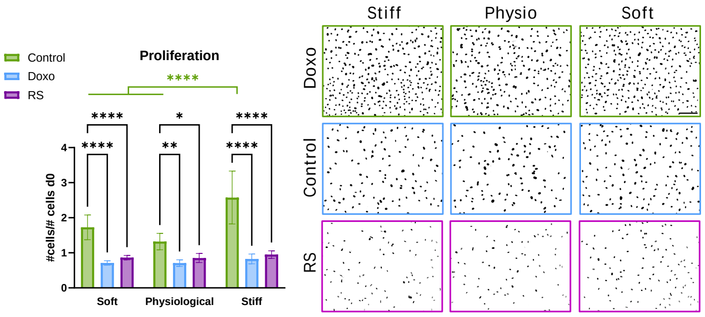


Figure S2: **Substrates' mechanical properties effect on proliferation** Green=Control, Blue=Doxo, Pink=RS. Left: Proliferation ratio is defined as the number of cells at day 7 over the number of cells at day 0 (mean ± SD). Statistical analysis: two-way Anova, Tukey's comparison. N=2, n=3. Right: Representative images of cells stained with live-Hoechst. Images acquired with M700 EVOS Microscope, objective 10x. Scale bar: 250 $\mu$m.

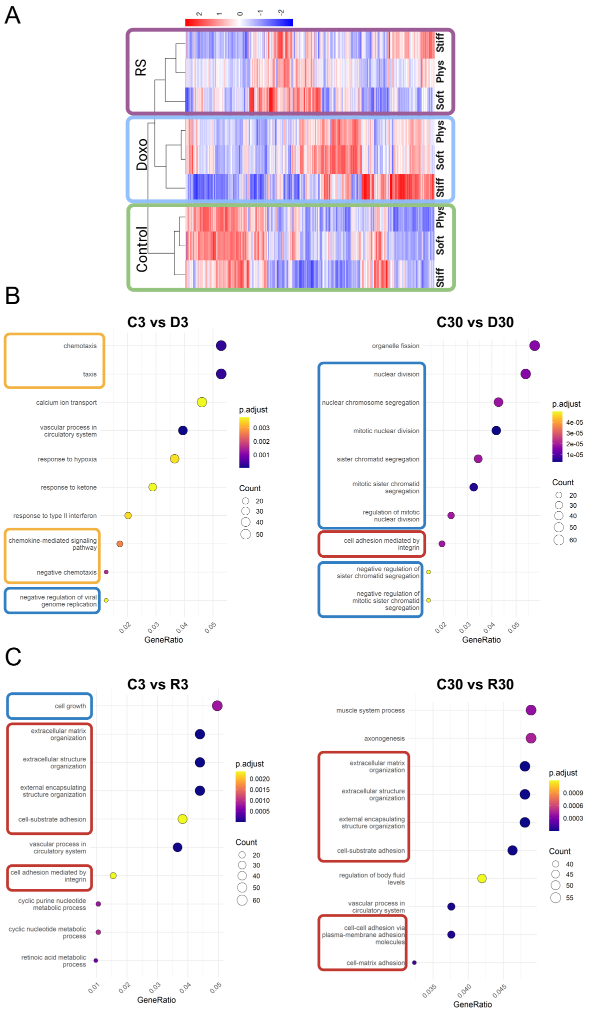


Figure S3: **RNA seq analysis results** (A) Heatmap obtained filtering the TPM data selecting the genes with TPM > 100 for at least one sample. 1003 genes are represented here. The clusters are indicated at the top of the graph. (B and C) Gene Ontology (GO) enrichment analysis displaying the top 10 Biological Process (BP) terms with highest statistical significance Control (C) and Doxo-treated cells (D) or Replicative senescent cells (R) on the soft (3kPa) and physiological (30kPa) substrates. For all GO term graphs, vertical axis shows GO terms and horizontal axis shows GeneRatio which corresponds to the genes of interest in the sample gene set divided over the total number of genes in the GO term. The size of each term dot represents the total count of genes identified in the sample gene set belonging to the term and the colour describes statistical significance according to the p-adjust legend. Recurring terms include chemotaxis, leukocyte migration, and extracellular matrix organization. Statistical analysis: DESeq2 (RNA-seq). N=3, n=3.
